# Automated model discovery for muscle using constitutive recurrent neural networks

**DOI:** 10.1101/2023.05.09.540027

**Authors:** Lucy M. Wang, Kevin Linka, Ellen Kuhl

**Affiliations:** Department of Mechanical Engineering, Stanford University, Stanford, California, United States

**Keywords:** automated model discovery, hyperelasticity, viscoelasticity, constitutive neural networks, recurrent neural networks, skeletal muscle

## Abstract

The stiffness of soft biological tissues not only depends on the applied deformation, but also on the deformation rate. To model this type of behavior, traditional approaches select a specific time-dependent constitutive model and fit its parameters to experimental data. Instead, a new trend now suggests a machine-learning based approach that simultaneously discovers both the best model and best parameters to explain given data. Recent studies have shown that feed-forward constitutive neural networks can robustly discover constitutive models and parameters for hyperelastic materials. However, feed-forward architectures fail to capture the history dependence of viscoelastic soft tissues. Here we combine a feed-forward constitutive neural network for the hyperelastic response and a recurrent neural network for the viscous response inspired by the theory of quasi-linear viscoelasticity. Our novel rheologically-informed network architecture discovers the time-independent initial stress using the feed-forward network and the time-dependent relaxation using the recurrent network. We train and test our combined network using unconfined compression relaxation experiments of passive skeletal muscle and compare our discovered model to a neo Hookean standard linear solid and to a vanilla recurrent neural network with no mechanics knowledge. We demonstrate that, for limited experimental data, our new constitutive recurrent neural network discovers models and parameters that satisfy basic physical principles and generalize well to unseen data. We discover a Mooney-Rivlin type two-term initial stored energy function that is linear in the first invariant *I*_1_ and quadratic in the second invariant *I*_2_ with stiffness parameters of 0.60kPa and 0.55kPa. We also discover a Prony-series type relaxation function with time constants of 0.362s, 2.54s, and 52.0s with coefficients of 0.89, 0.05, and 0.03. Our newly discovered model outperforms both the neo Hookean standard linear solid and the vanilla recurrent neural network in terms of prediction accuracy on unseen data. Our results suggest that constitutive recurrent neural networks can autonomously discover both model and parameters that best explain experimental data of soft viscoelastic tissues. Our source code, data, and examples are available at https://github.com/LivingMatterLab.

## 1 Introduction

The mechanical behavior of biological tissues exhibits many complexities that need to be taken into consideration in constitutive modeling [21]. For example, we can observe nonlinearity [42], heterogeneity [8], anisotropy [36], and viscoelasticity [12] in a variety of tissues ranging from tendon [37] to liver [42]. As an alternative to traditional constitutive modeling, researchers are now exploring the ability of neural networks to capture the intricate mechanical response of various biological tissues [35] including skin [29, 52], arteries [23], and the brain [32, 34, 51]. Traditional constitutive modeling assumes a certain functional form, and fits the parameters to the measured data. This may introduce significant modeling errors if the assumed functional form does not represent the material behavior well. In contrast, neural networks have the potential to learn both the functional form and its parameters [3, 24, 49], creating a more accurate representation of the material behavior.

A classical feed-forward neural network can capture traits such as nonlinearity [32, 34], heterogeneity [51], and anisotropy [23, 29], but these architectures are not well suited for modeling viscoelasticity. Instead, recurrent neural network architectures can model history-dependent behaviors such as viscoelasticity, where information from previous time steps informs the material response at the current time point [6]. In the classical deep learning realm, recurrent neural networks are successfully applied to sequence problems [39] like natural language processing [15], signal processing [71], and robot control [68]. In constitutive modeling, researchers are now beginning to evaluate the potential of recurrent neural networks to model plasticity [7, 54], viscoelasticity [1, 10, 43], and fatigue damage [67]. These recurrent neural networks also include various approaches such as directly predicting stress from strain input [10] and incorporating the recurrent neural network as a surrogate for micro level response in multiscale simulations [14].

In the early applications of recurrent neural networks to constitutive modeling, researchers directly adopted recurrent network architectures from the classical deep learning field [10,17,54,70]. While these classical recurrent network architectures reproduce history-dependent material behaviors, they typically involve hundreds to thousands of parameters, if not more. As a result, classical recurrent networks require large amounts of training data which may not be practical when building a constitutive model based on limited experimental measurements [2]. Furthermore, the parameters in these classical networks have no clear physical interpretation, and their predictions may violate physical laws and constraints [31].

To address these concerns, researchers have proposed physics-informed neural networks that incorporate physics knowledge into neural network design [11, 69]. Integrating our prior physics knowledge reduces the amount of required training data and constrains solutions to a physically admissible subspace. Two conceptually different approaches have emerged to incorporate physics knowledge: The first approach adds physical constraints to the loss function to enforce thermodynamic principles [7, 33, 46]; the second approach hardwires physical constraints into the network input, architecture, and output to learn stored energy functions and evolution laws for internal state variables [19, 30, 31, 38]. The stress then follows from these intermediate functions by direct calculation of mechanics equations.

Here we propose a new physics-informed approach that combines a feed-forward hyperelastic neural network [31, 51] with a recurrent linear viscous neural network to model the viscoelastic behavior of soft biological tissues. To motivate our new network architecture, we briefly revisit the theory of quasi-linear viscoelasticity [13] in Section 2, and illustrate how it decomposes the total stress response into a time-independent initial elastic stress and a time-dependent viscous overstress. In Section 3, we map this theory onto a new network architecture that integrates a feed-forward hyperelastic network and a recurrent viscous network. We illustrate the modular nature of this approach by probing two alternative hyperelastic networks: principal-stretch-based [51] and invariant-based [31]. In Section 4, we test and train both networks on five unconfined compression relaxation tests of passive skeletal muscle [59], and compare both against an overly constrained model, the neo Hookean standard linear solid, and an overly flexible model, a vanilla recurrent neural network. We perform two separate tasks, train on one and test on four versus train on four and test on one, and demonstrate that our new networks can uniquely discover model and parameters that best explain the experimental data. We close with some limitations of our study in Section 5 and with a brief conclusion in Section 6.

## 2 Theory of quasi-linear viscoelasticity

We begin by revisiting the theory of quasi-linear viscoelasticity, first in three dimensions, and then for the special case of uniaxial tension and compression. To characterize the deformation of the sample we want to test, we introduce the deformation map ***φ*** that maps material particles ***X*** from the undeformed configuration to particles, ***x*** = ***φ***(***X***), in the deformed configuration. We describe relative deformations within the sample using the deformation gradient ***F***, the gradient of the deformation map ***φ*** with respect to the undeformed coordinates ***X*** and its Jacobian *J*,

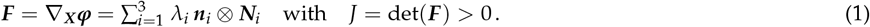

The spectral representation introduces the principal stretches *λ*_*i*_ and the principal directions ***N***_*i*_ and ***n***_*i*_ in the undeformed and deformed configurations, where ***F · N***_*i*_ = *λ*_*i*_***n***_*i*_. We further introduce the left Cauchy Green deformation tensor,

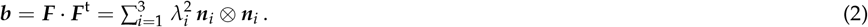

To characterize the isotropic material behavior, we introduce the three principal invariants, *I*_1_, *I*_2_, *I*_3_, either in terms of the principal stretches *λ*_1_, *λ*_2_, *λ*_3_,

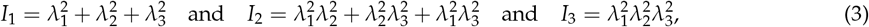

or in terms of the deformation gradient ***F***, with derivatives, *∂*_***F***_ *I*_1_, *∂*_***F***_ *I*_2_, *∂*_***F***_ *I*_3_,

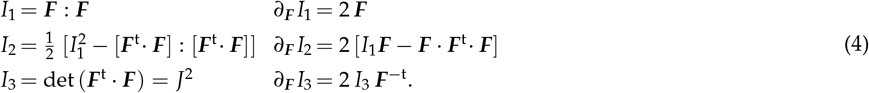

For isotropic, perfectly incompressible materials, the third invariant always remains identical to one, 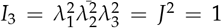, and the set of invariants reduces to *I*_1_ and *I*_2_. Following standard arguments of thermodynamics, we introduce an instantaneous elastic free energy function *ψ*_0_(***F***) from which we derive the instantaneous elastic Cauchy stress ***σ***_0_,

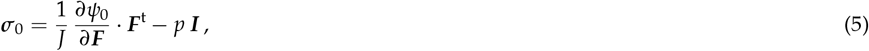

where *p* is the hydrostatic pressure that we determine from the boundary conditions and ***I*** is the second-order unit tensor. For an initial free energy expressed in terms of the principal stretches, *ψ*_0_(*λ*_1_, *λ*_2_, *λ*_3_), the initial Cauchy stress takes the following explicit representation,

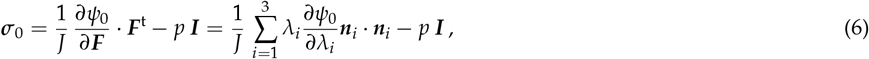

whereas for an energy function in terms of the invariants *ψ*_0_(*I*_1_, *I*_2_), the initial Cauchy stress takes the following form,

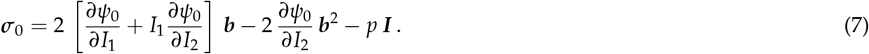

We pull the Cauchy stress back onto the undeformed reference configuration to obtain the initial Piola Kirchhoff stress,

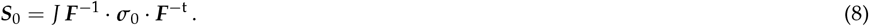

Following the quasi-linear viscoelastic theory [13], we introduce the viscoelastic Piola Kirchhoff stress through the following convolution integral,

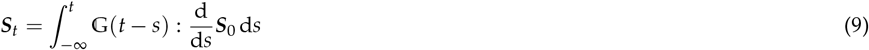

where 𝔾(*t*) is the time-dependent fourth-order viscous relaxation tensor and d***S***_0_/d*s* is the material time derivative of the instantaneous elastic Piola Kirchhoff stress according to equation (8). We assume that the relaxation is isotropic, and express it in terms of a discrete Prony series,

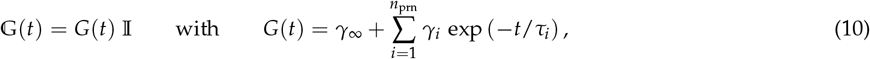

where *G*(*t*) is the time-dependent viscous relaxation function, 𝕀 is the fourth-order unit tensor, *γ*_∞_ and *γ*_*i*_ are the long-term modulus and the viscous relaxation coefficients with *γ*_∞_ + ∑_*i*=1_ *γ*_*i*_ = 1 and 0 ≤ *γ*_∞_, *γ*_*i*_ ≤ 1, *τ*_*i*_ are the viscous relaxation times of the *i* = 1, …, *n*_prn_ Maxwell elements, with *τ*_*i*_ > 0. We substitute the relaxation function (10) into the total stress expression (9), and push it forward into the deformed current configuration to obtain the total Cauchy stress,

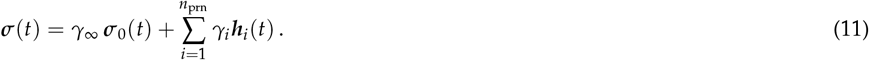

Here we have introduced a set of stress-like variables, the tensor-valued spatial overstresses ***h***_*i*_ for each Maxwell element *i*, as a push-forward of their material counterparts,

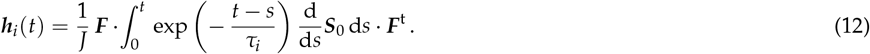

Since the exponential function gradually decays to zero, the viscous overstresses in equation (12) gradually decay in time, ***h***_*i*_(*t*→ ∞) = **0**. The total stress, ***σ***, in equation (11) converges towards the long-term equilibrium stress, ***σ***_∞_ = *γ*_∞_ ***σ***_0_.

### Uniaxial tension and compression

For the special case of uniaxial tension and compression, we deform the specimen in one direction, *F*_11_ = *λ*_1_ = *λ*, where the stretch *λ* = *l*/*L* denotes the relative change in length. For an isotropic, perfectly incompressible material with 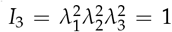, the stretches orthogonal to the loading direction are identical and equal to the inverse of the square root of the stretch, *F*_22_ = *λ*_2_ = *λ*^−1/2^ and *F*_33_ = *λ*_3_ = *λ*^−1/2^. From the resulting deformation gradient,

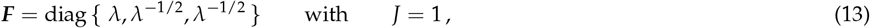

and left Cauchy Green deformation tensor,

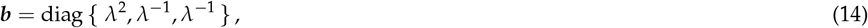

we calculate the first and second invariants and their derivatives,

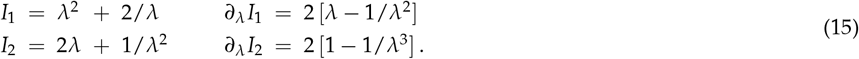

We derive the instantaneous elastic Cauchy stress *σ*_0_ = *σ*_11_ in the loading direction in terms of the instantaneous elastic free energy function *ψ*_0_ for perfectly incompressible materials using standard arguments of thermodynamics,

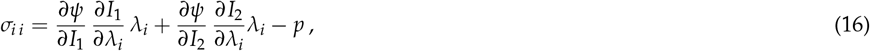

for *i* = 1, 2, 3. Here, *p* denotes the hydrostatic pressure that we determine from the zero stress condition in the transverse directions, *σ*_22_ = 0 and *σ*_33_ = 0, using equation (16) as *p* = [2/*λ*] *∂ψ*/*∂I*_1_ + [2*λ* + 2/*λ*^2^] *∂ψ*/*∂I*_2_. This results in the following instantaneous elastic uniaxial stress-stretch relation for perfectly incompressible, isotropic materials,

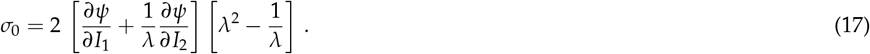

The underlying assumption of the theory of quasi-linear viscoelasticity is that we can multiplicatively decompose the total viscoelastic stress *σ*(*t*) into the strain-dependent function *σ*_0_ from equation (17) and a dimensionless time-dependent function *g*(*t*) [13]. We can then represent the total viscoelastic Cauchy stress through the convolution integral,

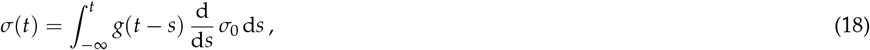

where *g*(*t*) is the time-dependent viscous relaxation function and d*σ*_0_/d*s* is the material time derivative of the instantaneous elastic uniaxial Cauchy stress according to equation (17). According to the theory of quasi-linear viscoelasticity and motivated by the generalized Maxwell model, we choose a discrete Prony series for the relaxation function,

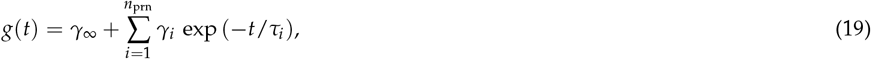

where *γ*_∞_ and *γ*_*i*_ are the long-term modulus and the viscous relaxation coefficients with *γ*_∞_ + ∑_*i*_ *γ*_*i*_ = 1 and 0 ≤ *γ*_∞_, ≤*γ*_*i*_ 1, *τ*_*i*_ are the viscous relaxation times of the *i* = 1, …, *n*_prn_ Maxwell elements, with *τ*_*i*_ *>* 0. By substituting equation (19) into equation (18) we obtain a convenient expression for the total Cauchy stress,

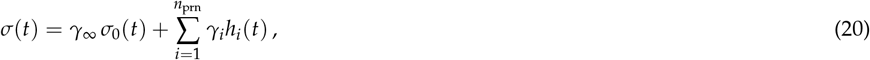

where we have introduced a set of new stress-like variables, the scalar-valued overstresses, *h*_*i*_(*t*), for each Maxwell element *i*,

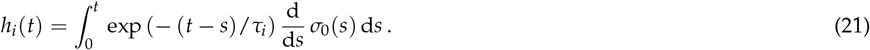

In general, the above convolution integral does not have a closed form solution, and we solve it numerically using an explicit Euler forward time integration scheme [55]. We discretize the time interval of interest and march forward from the previous time point *t*_*n*_ to the current time point *t*_*n*+1_ with the discrete time step size, Δ*t* = *t*_*n*+1_ − *t*_*n*_, using the following update equations for the total stress [22, 50],

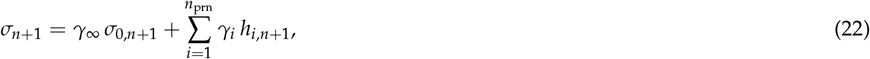

in terms of the *i* = 1, …, *n*_prn_ overstresses,

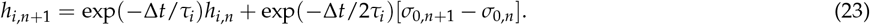

Before we move on to the next time step, we need to store both the total stress *σ*_*n*+1_ and the overstresses *h*_*i,n*+1_ of the current time step. Motivated by the considerations in this section, we will now introduce four different approaches to model the viscoelastic behavior of passive skeletal muscle.

## 3 Constitutive recurrent neural networks

In this section, we propose two constitutive recurrent neural networks inspired and informed by the quasi-linear viscoelastic theory in Section 2: one principal-stretch-based and one invariant-based. For comparison, we also introduce a neo Hookean standard linear solid, and a vanilla recurrent neural network. Figure 1 illustrates our constitutive recurrent neural network that combines a feed-forward network, top left and right, to discover the initial elastic stress and its parameters with a recurrent neural network, bottom, to discover the viscous overstress with its viscous parameters. We demonstrate both a principal-stretch-based network, top left, and a feed-forward invariant-based network, top right, for the initial elastic stress.

**Figure 1:**
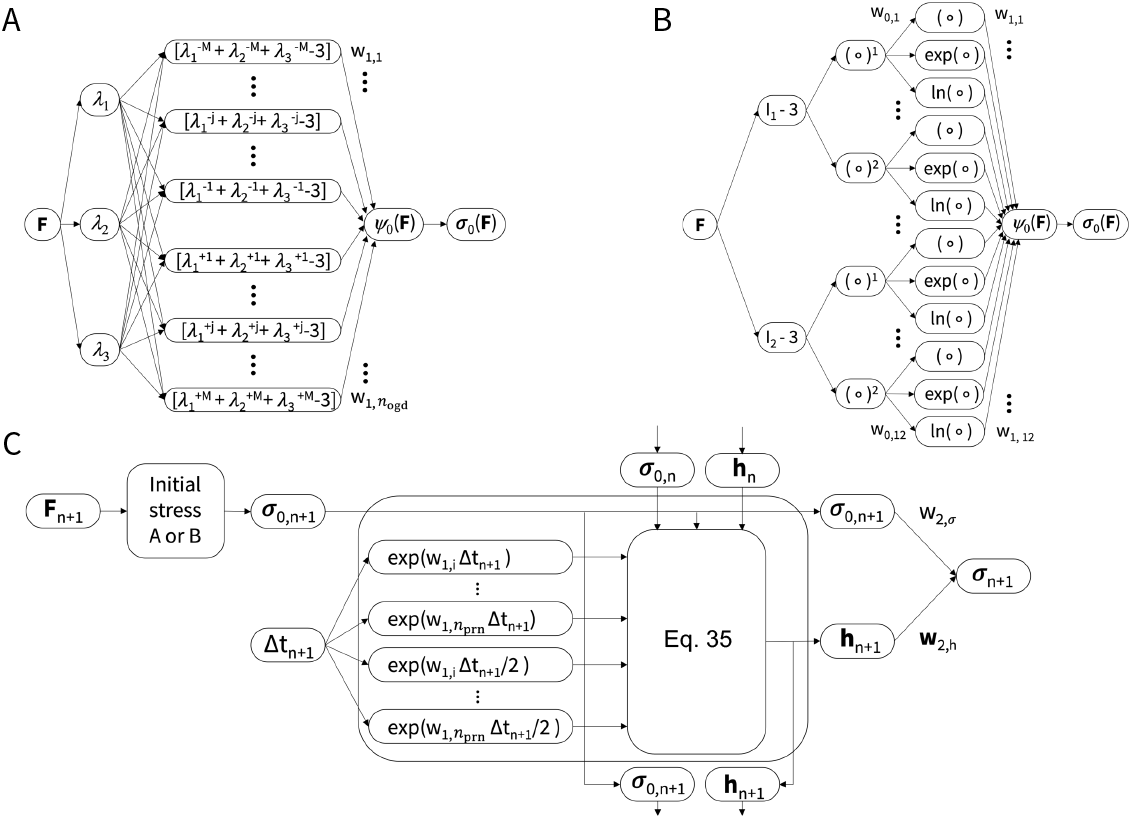
Principal-stretch-based and invariant-based recurrent neural network. The network takes the deformation gradient ***F***_*n*+1_ and time step size Δ*t*_*n*+1_ as input and outputs the total Cauchy stress ***σ***_*n*+1_. The network architecture combines a feed-forward constitutive neural network block to discover the time-independent initial stress function ***σ***_0,*n*+1_and its elastic parameters, followed by a recurrent neural network block to discover the time-dependent relaxation function and its Prony parameters. The network has a modular architecture and can seamlessly integrate different building blocks, for example, a principal-stretch or invariant-based neural network for the initial stress.

### Principal-stretch-based elastic stress

The principal-stretch-based network in Figure 1, top left, is parameterized in terms of the principal stretches, *λ*_1_, *λ*_2_, *λ*_3_, that act as input to a single dense layer [51]. Its initial stored energy function is inspired by the classical Ogden model [44] with *n*_ogd_ terms with fixed exponents,

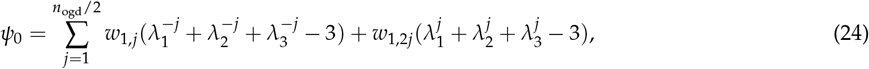

where *w*_1,*j*_ and *w*_1,2*j*_ are the weights learned by the network during training that we constrain to be positive, and 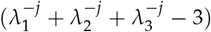 and 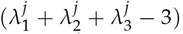, are the activation functions that raise the principal stretches to fixed powers 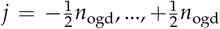. Unlike classical neural network architectures, the activation functions are applied first and the weights second. We can then derive the initial Cauchy stress as illustrated in Section 2 using equation (6) for the full stress tensor,

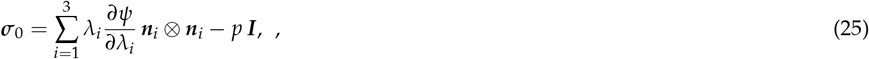

where *n*_*i*_ are the principal directions, or equation (16) for the initial uniaxial stress in the loading direction,

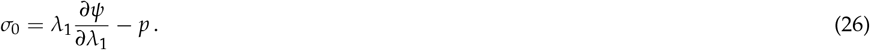

The hydrostatic pressure *p* follows from the zero-normal-stress condition as described in Section

### Invariant-based elastic stress

In contrast to the principal-stretch-based network, the invariant-based network in Figure 1, top right, is parameterized in terms of the first and second invariants, *I*_1_, *I*_2_, [34]. The network calculates both invariants using equation (4), and feeds them into two hidden layers with linear (○) and quadratic (○)^2^ activation functions in the first layer and linear (○), exponential (exp((○) 1)), and natural logarithmic ln(1 (○)) activation functions in the second layer. The resulting initial stored energy function has a total of *n*_inv_ = 12 terms,

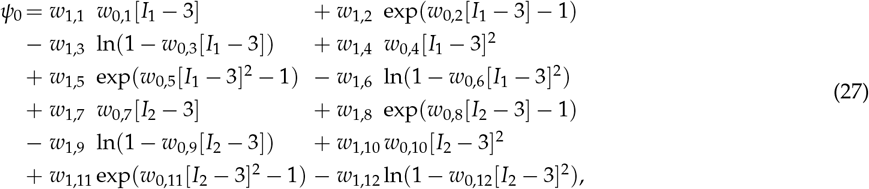

where *w*_0,1−12_ and *w*_1,1−12_ are the weights of the first and second layers, and are constrained to be positive. We can then derive the initial Cauchy stress as illustrated in Section 2 using equation (7) for the full stress tensor,

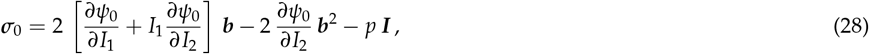

where the hydrostatic pressure *p* follows from the zero-normal-stress condition, or using equation (17) for the initial uniaxial stress in the loading direction,

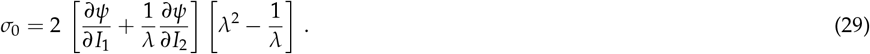

### Time-dependent viscous overstress

Figure 1, bottom, shows the recurrent neural network inspired by the relaxation function of the Prony series of equation (19),

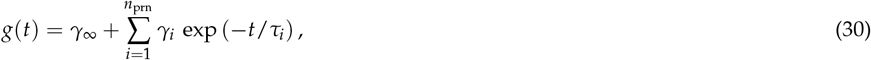

from which we derive the scalar-valued overstresses *h*_*i,n*+1_ according to equation (21),

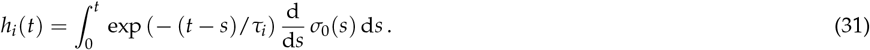

### Time-discrete update equations

Following Section (2), we obtain the time-discrete updates of the total stress according to equation (22),

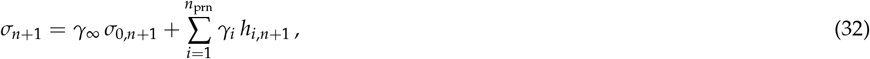

and of the overstresses according to equation (23),

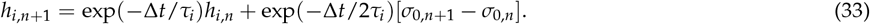

The architecture of our custom recurrent neural network in Figure 1, bottom, is inspired by these two update formulas. Its inputs are the initial stress *σ*_0_, either from equation (26) for the principal-stretch-based neural network, or from equation (29) for the invariant-based neural network, and the time step size Δ*t*. The recurrent neural network first calculates two sets of intermediate terms, *a*_*i*_ and *b*_*i*_, from the time step size, Δ*t*_*n*+1_,

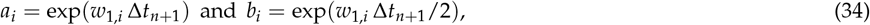

where *w*_1,*i*_ are the *i* = 1, …, *n*_prn_ weights that the network learns during training. Upon comparison with equation (33), we see that the network weights are equal to the negative inverse relaxation times, *w*_1,*i*_ = − 1/*τ*_*i*_. To ensure that the time constants are positive, we constrain the weights to be negative. After this initial layer, the network updates the overstresses *h*_*i*_ using equation (33),

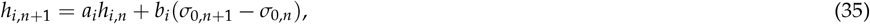

where *a*_*i*_ and *b*_*i*_ are the intermediate terms from equations (34), *σ*_0,*n*+1_ is the initial stress output from the principal-stretch-based or invariant-based neural network, and *σ*_0,*n*_ and *h*_*i,n*_ are passed forward from the previous time step *t*_*n*_. The final layer is a dense layer that updates the total Cauchy stress according to equation (32),

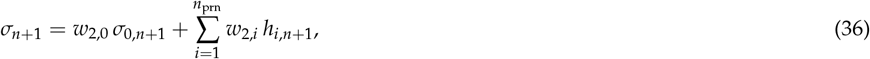

where *w*_2,0_ and *w*_2,*i*_ are the weights learned by the model during training. Upon comparison to equation (32), we see that the weights are equal to the long-term modulus, *w*_2,0_ = *γ*_∞_, and the viscous relaxation coefficients, *w*_2,*i*_ = *γ*_*i*_. We constrain the weights to be positive and to sum to one in accordance with *γ*_∞_ + ∑_*i*_ *γ*_*i*_ = 1.

### Loss function

We select a principal-stretch-based network with *n*_ogd_ = 20 Ogden terms in equation (24) and *n*_prn_ = 10 Prony series terms in equation (36). For these values, the principal-stretch-based network has a total of 41 trainable parameters or weights, 20 for the principal stretch terms 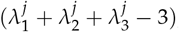, ten for the negative inverse relaxation times −1/*τ*_*i*_, one for the initial stress *σ*_0,*n*+1_, and ten for the overstresses *h*_*i,n*+1_. We select an invariant-based network with *n*_inv_ = 12 invariant terms in equation (27) and *n*_prn_ = 10 Prony series terms in equation (30). For these values, the invariant-based network has a total of 45 trainable parameters or weights, two times twelve for the invariant terms, ten for the negative inverse relaxation times −1/*τ*_*i*_, one for the initial stress *σ*_0,*n*+1_, and ten for the overstresses *h*_*i,n*+1_. We learn these network weights ***θ*** = {***w***} by minimizing a loss function of mean squared error type, parameterized in terms of the stretch *λ* and time *t*,

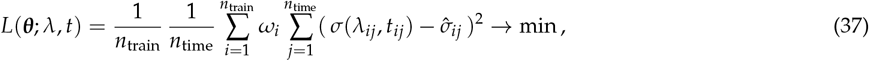

where *i* = 1, …, *n*_train_ denotes the number of training sets, *j* = 1, …, *n*_time_ denotes the number of data points in the time series, and 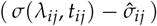 is the difference between the model stress *σ*(*λ*_*ij*_, *t*_*ij*_) at stretch *λ*_*ij*_ and time *t*_*ij*_ and the experimental stress 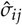. We weight the loss associated with each training set with the coefficient, 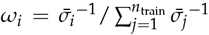 to normalize the contributions from each of the training sets since they have different mean stress values 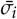. We train both networks by minimizing the loss function for 5000 epochs using the Adam optimizer, with a learning rate *α* = 0.001 and parameters *β*_1_ = 0.9 and *β*_2_ = 0.999.

### Regularization

To investigate the effect of L2 regularization on model discovery, we add a penalty term to the loss function in equation (37) to penalize network weights with large magnitudes. We assess the effect of L2 regularization in both the feed-forward network and the recurrent neural network. With regularization in the feed-forward network, the loss function becomes

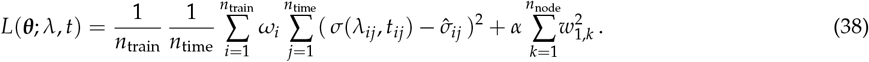

Here, *α* is the regularization parameter and *w*_1,*k*_ are the weights of the last hidden layer of the feed-forward network with either *n*_node_ = *n*_ogd_ = 20 or *n*_node_ = *n*_inv_ = 12. We compare the effect of varying regularization parameters, *α*, ranging from 10^−7^ to 10^−1^ on the invariant-based network. With regularization in the recurrent network, the loss function becomes

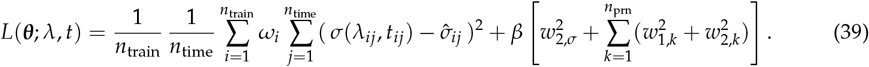

Here, *β* is the regularization parameter and *w*_2,*σ*_, *w*_1,*k*_, and *w*_2,*k*_ are the weights of the recurrent neural network. We compare the effect of varying *β* from 10^−4^ to 10^−1^.

### Special case: Neo Hookean standard linear solid

A special case of both the principal-stretch and invariant-based recurrent neural networks is the neo Hookean standard linear solid. It is the simplest of all physics-informed models, and also the most frequently used. We use this model as an overly-constrained, low-end comparison and expect that it will have difficulties fitting the nonlinear elastic behavior, but will not display non-physical responses for small data. The initial stored energy function of the neo Hookean standard linear solid is linear in the sum of the three principal stretches, *λ*_1_, *λ*_2_, *λ*_3_, squared, or in other words, linear in the first invariant *I*_1_,

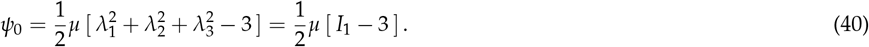

The shear modulus *μ* follows from the weights of the principal-stretch-based network as *μ* = 2*w*_1,12_ and from the weights of the invariant-based network as *μ* = 2*w*_0,1_*w*_1,1_ with all other weights identical to zero. For the special case of uniaxial tension and compression, the instantaneous elastic uniaxial stress-stretch relation for a perfectly incompressible, isotropic material of equation (17) becomes

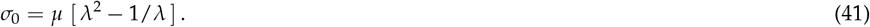

For the neo Hookean standard linear solid, the Prony series only has a single one term, *n*_*prn*_ = 1. To numerically solve for its total stress, we discretize the time interval of interest and adopt an explicit Euler forward scheme to march from time *t*_*n*_ to *t*_*n*+1_ in increments of Δ*t* = *t*_*n*+1_ − *t*_*n*_ using the update rules according to equation (22) for the total stress,

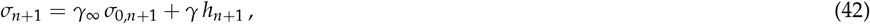

and equation (23) for the viscous overstress,

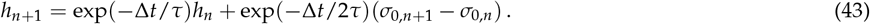

The classical neo Hookean standard linear solid has four parameters that we need to fit to the data: the shear modulus *μ* to characterize the instantaneous elastic behavior, and the long-term modulus *γ*_∞_, the relaxation coefficient *γ*, and the relaxation time *τ* to characterize the time-dependent viscous behavior.

### General case: Vanilla recurrent neural network

Recurrent neural networks have a built-in memory structure and learn functions that, at any given time point, depend on all past inputs. Here we adopt a vanilla recurrent neural network as an overly-flexible, high-end comparison and expect that it will not have difficulties fitting the nonlinear elastic behavior, but might display overfitting and non-physical responses for small data. For this vanilla type recurrent neural network, the new stress state *σ*_*n*+1_ at time *t*_*n*+1_ not only depends on the stretch and the new time point {*λ*_*n*+1_, *t*_*n*+1_}, but also on a history vector *h*_*n*_ from the previous time point *t*_*n*_ [16]. At each new time point, the network updates the history vector,

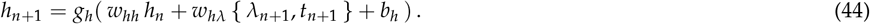

and calculates the new stress state using an additional feed-forward layer,

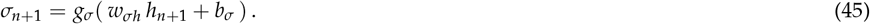

Importantly, a recurrent neural network learns the same weights *w*_*hh*_, *w*_*hλ*_, *w*_*σh*_ and biases *b*_*h*_, *b*_*σ*_ for all time steps, so equations (44) and (45) closely mirror the update rules for the viscous overstress (23) and the elastic stress (22) of the theory of quasi-linear viscoelasticity in Section 2. Here we choose a hyperbolic tangent activation function *g*_*h*_(○) for the history vector (44), a linear activation function *g*_*σ*_(○) for the stress (45). Our network input {*λ, t*} is a two-unit vector of stretch and time and our network output *σ* is a scalar. For the history vector *h*, we select an 8-unit vector, such that *w*_*hλ*_ is an 8×2 matrix, *w*_*hh*_ is an 8×8 matrix, *w*_*σh*_ is a 1×8 vector, *b*_*h*_ is an 8 ×1 vector, and *b*_*σ*_ is a scalar. This results in 16 + 64 + 8 + 8 + 1 = 97 total trainable parameters, ***θ*** = {***w, b***}. The network learns its parameters by minimizing the loss function that penalizes the error between model and data. We adopt a logarithmic hyperbolic cosine loss function parameterized in terms of the stretch *λ* and time *t*,

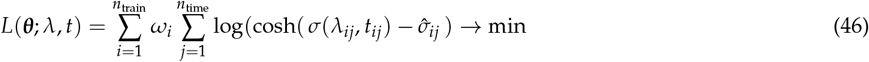

where *i* = 1, …, *n*_train_ denotes the number of training sets, *j* = 1, …, *n*_time_ denotes the number of data points in the time series, and 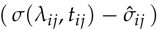 is the difference between the model stress *σ*(*λ*_*ij*_, *t*_*ij*_) at time *t*_*ij*_ and the experimental stress 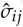. We normalize the loss associated with each training set with the weight, 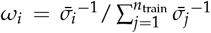 since each training set has a different mean stress values 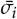. We train the model for 5000 epochs using the Adam optimizer, with a learning rate *α* = 0.1 and parameters *β*_1_ = 0.9 and *β*_2_ = 0.999.

#### 3.1 Data

We train and test all four models on unconfined compression relaxation tests of passive skeletal muscle collected from porcine gluteus muscle tissue [59]. The data consist of recordings from five different experiments in which the muscle is compressed to three different stretch levels at three different stretch rates. After an initial ramp loading, the final stretch level is held constant for 300 seconds. For three experiments, the stretch rate is fixed at 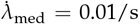, and the minimum compressive stretch is varied as *λ*_low_ = 0.9, *λ*_med_ = 0.8, and *λ*_high_ = 0.7. For three experiments, the minimum comprssive stretch is fixed at *λ*_high_ = 0.7, and the stretch rate is varied as 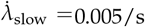, 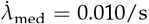 and 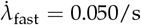. As the *λ*_high_ = 0.7, 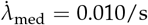 case is repeated, this results in a total of five data sets. To minimize the effects of tissue anisotropy all experiment are performed with compression in the muscle fiber direction. Table 1 summarizes the five data sets, reported as means over *n* = 6 tests per data set.

**Table 1:** Unconfined compression relaxation data of passive skeletal muscle. Compression of gluteus muscle samples at fixed stretch rate of 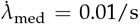 and varying stretches of *λ*_low_=0.9, *λ*_med_=0.8, and *λ*_high_=0.7, and at fixed stretch *λ*_high_ = 0.7 and varying stretch rates of 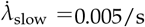, 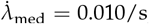, and 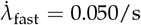. Cauchy stresses are reported as means from *n* = 6 samples [59].

| $\lambda_{\text{low}} = 0.9$<br>$\dot{\lambda}_{\text{med}} = 0.010/\text{s}$ | | | $\lambda_{\text{med}} = 0.8$<br>$\dot{\lambda}_{\text{med}} = 0.010/\text{s}$ | | | $\lambda_{\text{high}} = 0.7$<br>$\dot{\lambda}_{\text{med}} = 0.010/\text{s}$ | | | $\lambda_{\text{high}} = 0.7$<br>$\dot{\lambda}_{\text{slow}} = 0.005/\text{s}$ | | | $\lambda_{\text{high}} = 0.7$<br>$\dot{\lambda}_{\text{fast}} = 0.050/\text{s}$ | | |
| --- | --- | --- | --- | --- | --- | --- | --- | --- | --- | --- | --- | --- | --- | --- |
| $t$<br>[s] | $\lambda$<br>[-] | $\sigma$<br>[kPa] | $t$<br>[s] | $\lambda$<br>[-] | $\sigma$<br>[kPa] | $t$<br>[s] | $\lambda$<br>[-] | $\sigma$<br>[kPa] | $t$<br>[s] | $\lambda$<br>[-] | $\sigma$<br>[kPa] | $t$<br>[s] | $\lambda$<br>[-] | $\sigma$<br>[kPa] |
| 0.0 | 1.00 | 0.000 | 0.0 | 1.00 | 0.000 | 0.0 | 1.00 | 0.000 | 0.0 | 1.00 | 0.000 | 0.0 | 1.00 | 0.000 |
| 2.0 | 0.98 | -0.060 | 4.0 | 0.96 | -0.119 | 6.0 | 0.94 | -0.167 | 12.0 | 0.94 | -0.077 | 1.2 | 0.94 | -0.223 |
| 4.0 | 0.96 | -0.101 | 8.0 | 0.92 | -0.209 | 12.0 | 0.88 | -0.343 | 24.0 | 0.88 | -0.193 | 2.4 | 0.88 | -0.427 |
| 6.0 | 0.94 | -0.148 | 12.0 | 0.88 | -0.342 | 18.0 | 0.82 | -0.602 | 36.0 | 0.82 | -0.355 | 3.6 | 0.82 | -0.664 |
| 8.0 | 0.92 | -0.205 | 16.0 | 0.84 | -0.511 | 24.0 | 0.76 | -1.058 | 48.0 | 0.76 | -0.702 | 4.8 | 0.76 | -1.518 |
| 10.0 | 0.90 | -0.266 | 20.0 | 0.80 | -0.711 | 30.0 | 0.70 | -1.844 | 60.0 | 0.70 | -1.424 | 6.0 | 0.70 | -2.315 |
| 12.0 | 0.90 | -0.240 | 22.0 | 0.80 | -0.616 | 32.0 | 0.70 | -1.593 | 62.0 | 0.70 | -1.235 | 8.0 | 0.70 | -1.694 |
| 15.0 | 0.90 | -0.212 | 25.0 | 0.80 | -0.569 | 35.0 | 0.70 | -1.473 | 65.0 | 0.70 | -1.120 | 10.0 | 0.70 | -1.454 |
| 20.0 | 0.90 | -0.192 | 30.0 | 0.80 | -0.529 | 40.0 | 0.70 | -1.376 | 70.0 | 0.70 | -1.025 | 15.0 | 0.70 | -1.267 |
| 30.0 | 0.90 | -0.174 | 40.0 | 0.80 | -0.487 | 50.0 | 0.70 | -1.273 | 80.0 | 0.70 | -0.917 | 20.0 | 0.70 | -1.161 |
| 40.0 | 0.90 | -0.160 | 50.0 | 0.80 | -0.465 | 60.0 | 0.70 | -1.212 | 90.0 | 0.70 | -0.861 | 30.0 | 0.70 | -1.058 |
| 60.0 | 0.90 | -0.147 | 70.0 | 0.80 | -0.433 | 80.0 | 0.70 | -1.139 | 110.0 | 0.70 | -0.782 | 40.0 | 0.70 | -0.992 |
| 80.0 | 0.90 | -0.138 | 90.0 | 0.80 | -0.411 | 100.0 | 0.70 | -1.090 | 130.0 | 0.70 | -0.734 | 60.0 | 0.70 | -0.904 |
| 100.0 | 0.90 | -0.130 | 110.0 | 0.80 | -0.395 | 120.0 | 0.70 | -1.057 | 150.0 | 0.70 | -0.695 | 80.0 | 0.70 | -0.844 |
| 130.0 | 0.90 | -0.125 | 140.0 | 0.80 | -0.377 | 150.0 | 0.70 | -1.014 | 180.0 | 0.70 | -0.654 | 120.0 | 0.70 | -0.774 |
| 170.0 | 0.90 | -0.113 | 180.0 | 0.80 | -0.357 | 190.0 | 0.70 | -0.970 | 220.0 | 0.70 | -0.610 | 180.0 | 0.70 | -0.708 |
| 220.0 | 0.90 | -0.106 | 230.0 | 0.80 | -0.339 | 240.0 | 0.70 | -0.932 | 270.0 | 0.70 | -0.578 | 240.0 | 0.70 | -0.658 |
| 300.0 | 0.90 | -0.097 | 310.0 | 0.80 | -0.322 | 320.0 | 0.70 | -0.881 | 350.0 | 0.70 | -0.531 | 300.0 | 0.70 | -0.625 |

#### 3.2 Statistical analysis

We use two error measures to quantify the model performance during testing and training. The first error measure is the normalized root mean squared error, *NRMSE*,

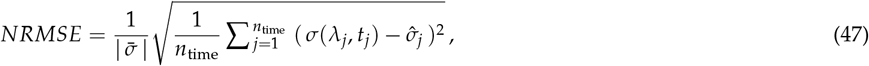

where 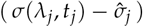 is the difference between the model stress *σ*(*λ*_*j*_, *t*_*j*_) and the experimental stress 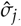, and *j* = 1, …, *n*_time_ denotes the number of data points in the time series. We normalize the error by the mean stress, 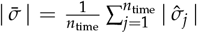 of each time series since the mean stress values of the five data curves in Section 3.1) vary by over an order of magnitude. The second error measure is the coefficient of determination, *R*^2^,

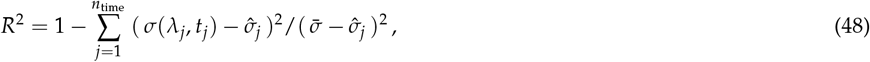

where 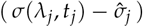 is the difference between the model stress *σ*(*λ*_*j*_, *t*_*j*_) and the experimental stress 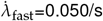, and 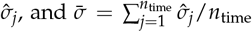 is the mean stress of the time series. In the discussion, we use the normalized root mean squared error to compare the different models because it provides more insight into the magnitude of the prediction errors compared to the magnitude of the data themselves. However, we also include the coefficient of determination to aid comparison to other studies. We performed all statistical analyses using the SciPy 1.7.3 Python library [60].

## 4 Results and Discussion

### 4.1 Recurrent neural network training

Figure 2 shows representative plots of the training and test losses for all three recurrent neural networks. The training loss of all three recurrent neural networks, vanilla, principal-stretch-based, and invariant based, converged within 5000 epochs. After confirming that 5000 epochs were generally sufficient for the training loss to plateau, we performed all following network training trials using 5000 epochs. The test loss of the principal-stretch and invariant-based networks displayed a similar tendency and dropped visibly within the first 2000 epochs. Notably, the test loss of the vanilla network remained unchanged across all epochs for the train on one case, and even increased after about 2000 epochs for the train on four case. This generalization gap indicates the poor predictive performance of the vanilla network when trained on limited data.

**Figure 2:**
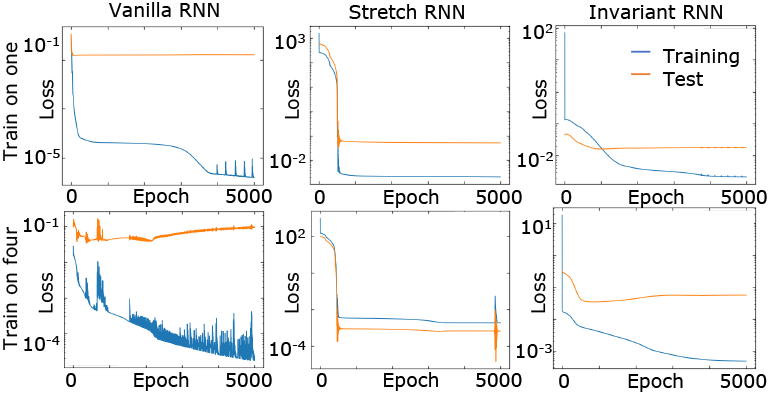
Representative evolution of training and test losses during model training. Columns display the three networks, the vanilla, principal-stretch-based, and invariant-based recurrent neural networks; rows represent the two tasks, train on one and test on the remaining four, and train on four and test on the remaining one.

### 4.2 Model performance

We evaluate all four models on two separate tasks, train on one data set and test on the remaining four; and train on four data sets and test on the remaining one. Figures 3 through 10 illustrate the models’ performance for both tasks. In these figures, each column represents one of five experiments with the corresponding column label signifying the training curve for the train on one task or the test curve for the train on four task. Each row represents a data curve, and the plots in that row display the model’s performance in fitting that data curve. If the curve was included in the training set, the annotation for the coefficient of determination is labeled 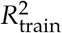, and if it was not included in the training set, the annotation is labeled 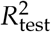.

**Figure 3:**
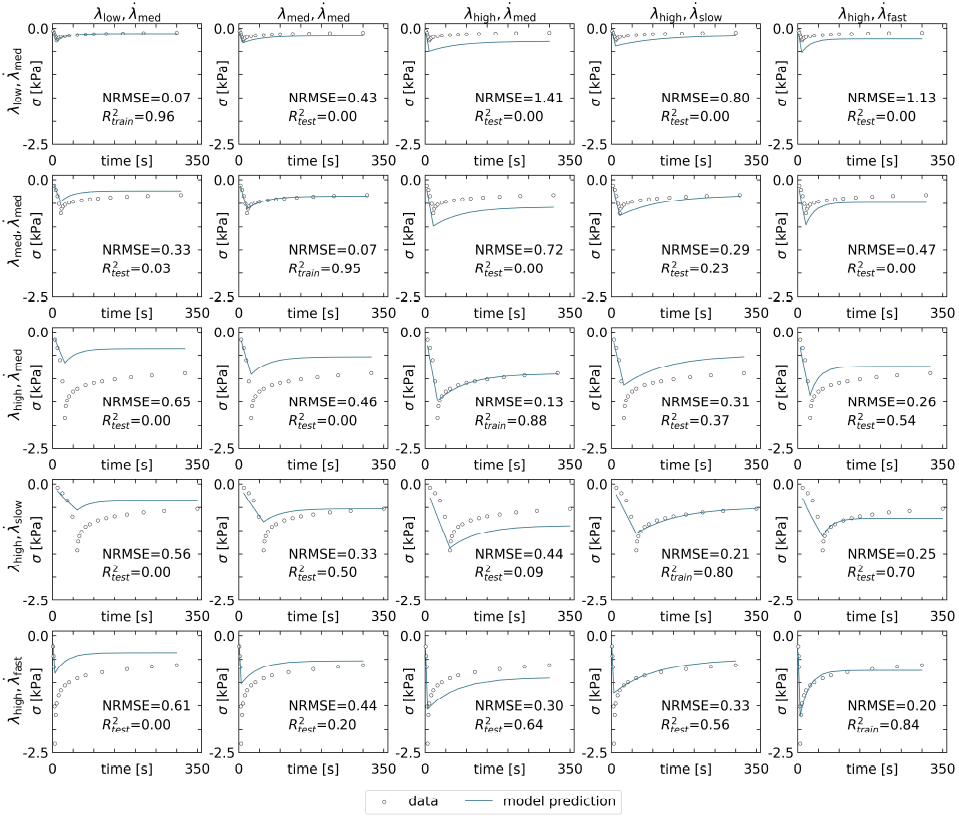
Performance of neo Hookean standard linear solid trained on one curve. Experimental data and discovered model for the constitutive behavior of muscle tissue after discovering the model parameters for just one of the data sets. Columns represent all five training runs with each column label signifying the training set. Rows display the model performance for all five data sets with the training cases on the diagonal and the test cases on the off-diagonal.

#### Train on one, test on four

In the first evaluation task, we trained the models on one of the five compression relaxation data sets from Table 1 and then tested the models’ predictive abilities on the remaining four curves. Figure 3 shows the results for the neo Hookean standard linear solid. As expected, since the model parameters are fit to the training curve, the errors are lowest for the training set (average NRMSE: 0.136) and higher for the test curves (average NRMSE: 0.526). The simplicity of the model makes it impossible to exactly match the shape of the experimental curves. This extreme case illustrates the limitations of building a constitutive model by first assuming a functional form and then fitting the parameters. The model fails to fit the experimental data for any combination of parameter values because it assumes that the stress will increase linearly, but this is not how the tissue behaves.

As an alternative to first assuming a functional form, vanilla recurrent neural networks learn both the functional form and its parameters from the data themselves. Figure 4 shows the performance of the vanilla recurrent neural network. With 97 trainable parameters, the vanilla recurrent neural network fits the training data sets almost perfectly (average NRMSE: 0.012), but is at risk for over-fitting to the training data. Overfitting can be observed clearly in the middle column of Figure 4 where the neural network predicts almost identical curves in the first three rows. The average errors on the test set (average NRMSE: 1.622) are much higher than those of the training set. Besides overfitting, the vanilla recurrent neural network sometimes produces predictions that violate physical principles. For example, the predicted stresses sometimes switch between increasing and decreasing during the ramp portion of the time history.

**Figure 4:**
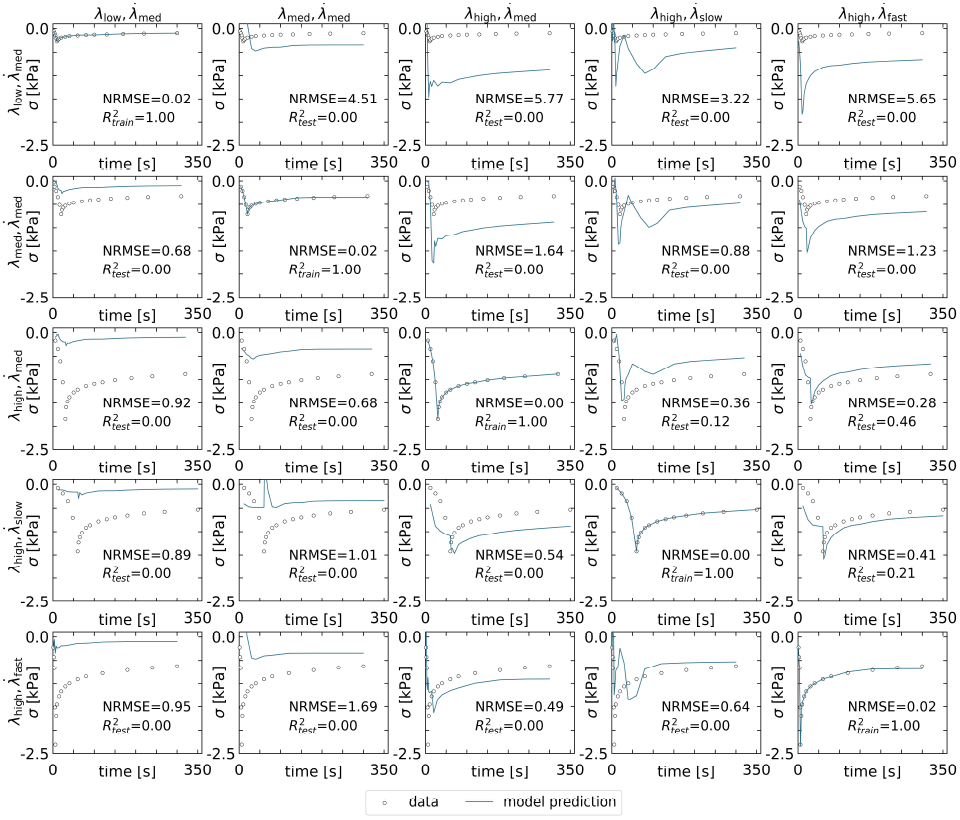
Performance of vanilla recurrent neural network trained on one curve. Experimental data and discovered model for the constitutive behavior of muscle tissue after discovering the model parameters for just one of the data sets. Columns represent all five training runs with each column label signifying the training set. Rows display the model performance for all five data sets with the training cases on the diagonal and the test cases on the off-diagonal.

Figures 5 and 6 show the results for the two constitutive recurrent neural network models. The performance of both the principal stretch version in Figure 5 and the invariant version in Figure 6 falls between that of the neo Hookean standard linear solid and the vanilla recurrent neural network when looking at the training sets (average NRMSE: 0.048 and 0.022). Both constitutive recurrent neural networks discover a functional form from the data and can approximate the data more closely compared to the restrictive neo Hookean standard linear solid model. However, the functional forms of both constitutive networks are more limited in scope than the vanilla network. Therefore, the constitutive networks cannot fit the training data as perfectly as the vanilla network. This limitation on the discovered functional forms provides a benefit, however, when looking at the test data sets. Compared to the vanilla network, both constitutive networks exhibit less overfitting as evidenced by the lower test errors (average NRMSE: 0.289 and 0.243). Importantly, unlike the vanilla network, the constitutive recurrent neural networks do not produce unphysical solutions with spurious oscillatory stress responses.

**Figure 5:**
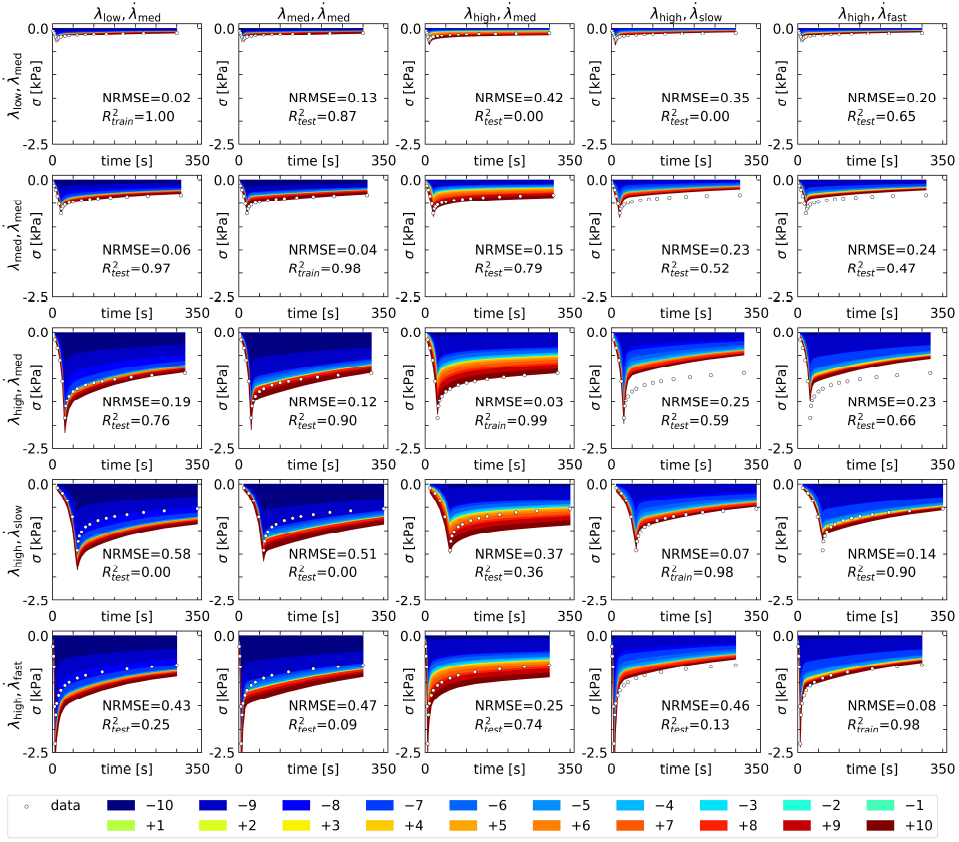
Performance of principal-stretch-based recurrent neural network trained on one curve. Experimental data and discovered model for the constitutive behavior of muscle tissue after discovering the model parameters for just one of the data sets. Columns represent all five training runs with each column label signifying the training set. Rows display the model performance for all five data sets with the training cases on the diagonal and the test cases on the off-diagonal. The contributions of the principal stretch terms in the initial stored energy function are illustrated in colors.

**Figure 6:**
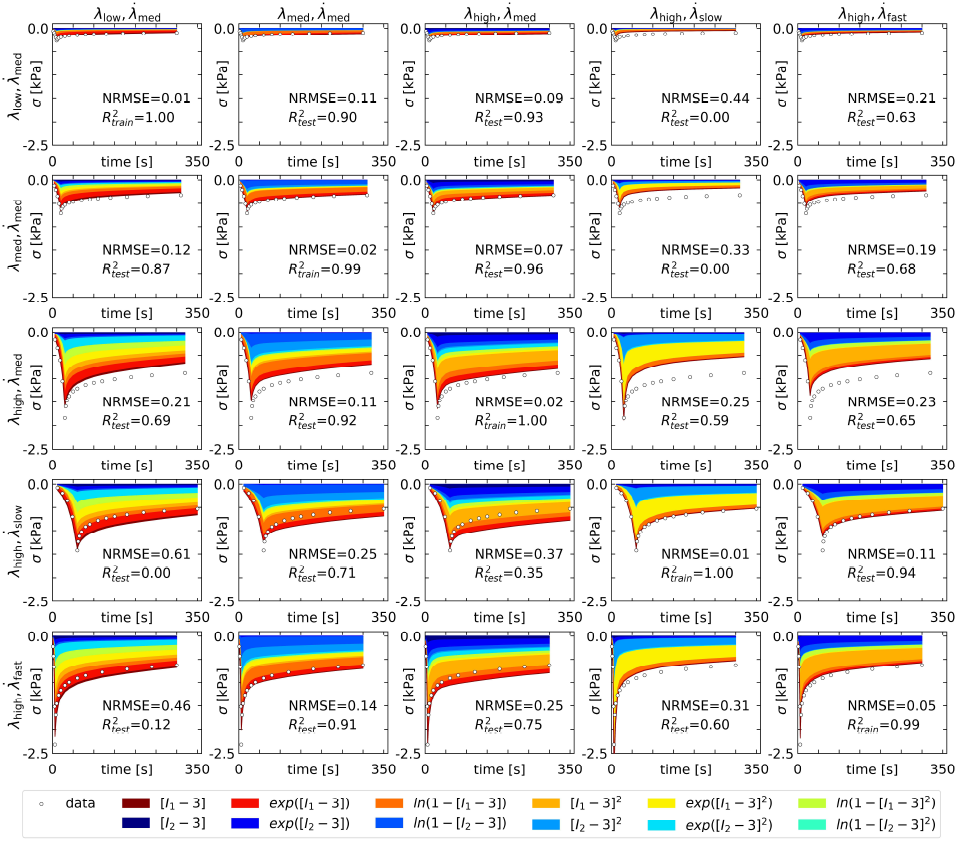
Performance of invariant-based recurrent neural network trained on one curve. Experimental data and discovered model for the constitutive behavior of muscle tissue after discovering the model parameters for just one of the data sets. Columns represent all five training runs with each column label signifying the training set. Rows display the model performance for all five data sets with the training cases on the diagonal and the test cases on the off-diagonal. The contributions of the principal stretch terms in the initial stored energy function are illustrated in colors.

#### Train on four, test on one

In the second evaluation task, we trained the four models on four out of five data curves and tested the models’ predictive abilities on the final remaining curve. Figure 7 shows the results for the neo Hookean standard linear solid model. Compared to the train on one task, the errors on the training set are higher for the train on four task, (increasing average NRMSE: from 0.136 to 0.370). The model struggles to find a single set of parameters that can describe four different curves simultaneously. With the increase in training data, however, the error on the test set becomes smaller (decreasing average NRMSE: from 0.526 to 0.462). Similar to the train on one task, the neo Hookean standard linear solid continues to be restricted by its simple functional form. It simply cannot fit the data well because the tissue does not behave in accordance with the assumed functional form.

**Figure 7:**
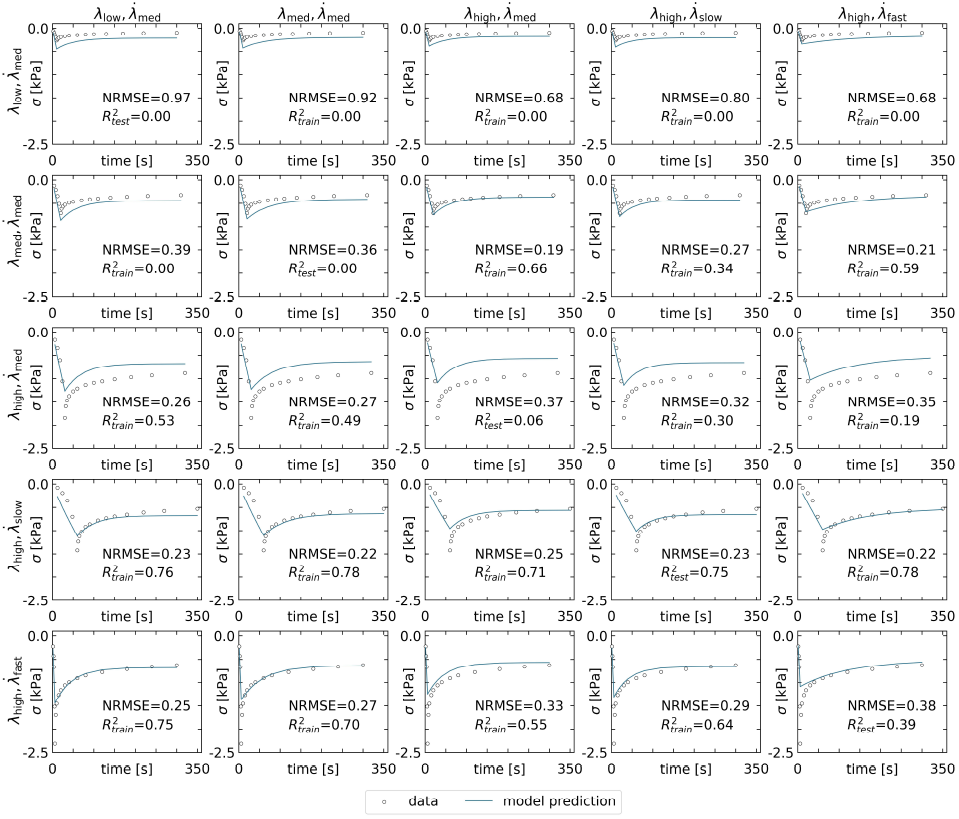
Performance of neo Hookean standard linear solid trained on four curves. Experimental data and discovered model for the constitutive behavior of muscle tissue after discovering the model parameters for four of the data sets combined. Columns represent all five training runs with each column label signifying the test set. Rows display the model performance for all five data sets with the training cases on the off-diagonal and the test cases on the diagonal.

Figure 8 shows that the vanilla recurrent neural network almost perfectly fits the training data set, similar to the train on one task but with a slightly larger error (increasing average NRMSE: from 0.012 to 0.040). In contrast to the neo Hookean standard linear solid, the vanilla recurrent neural network has the ability to learn a complex enough functional form to simultaneously fit four different curves using the same set of parameters. With the additional training data, the average test error goes down (decreasing average NRMSE: from 1.622 to 0.838). However, the issue of over-fitting remains apparent in comparing the train and test errors (average NRMSE: 0.040 vs. 0.838). Additionally, the issue of unphysical predicted solutions remains as seen clearly in the bottom left subplot of Figure 8. The vanilla network prediction in this test case shows the stress decreasing and then increasing all during the hold portion of the experiment where the stress is expected to monotonically decrease. These observations suggest that although vanilla recurrent neural network architectures can successfully learn history-dependent constitutive laws [10, 17, 54, 70], the amount of required training data may not be practical to collect in a typical experimental setup. Importantly, in this case, increasing the amount of data does not imply increasing the number of data points on a curve but rather increasing the number of different curves from varying experiments.

**Figure 8:**
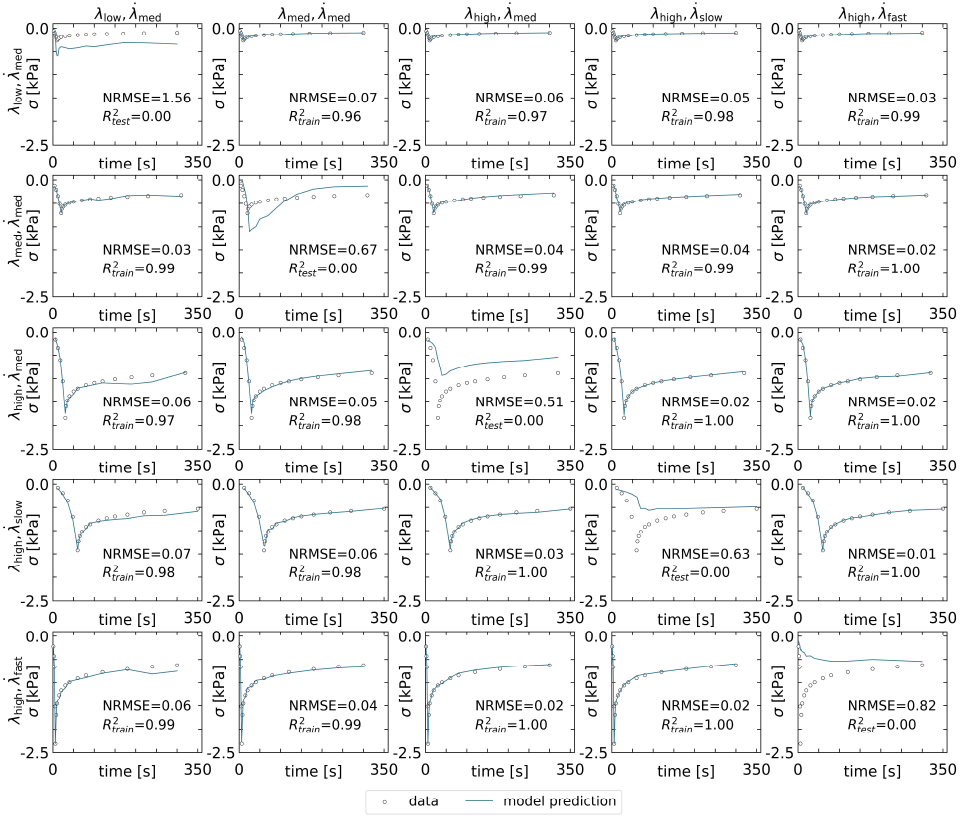
Performance of vanilla recurrent neural network trained on four curves. Experimental data and discovered model for the constitutive behavior of muscle tissue after discovering the model parameters for four of the data sets combined. Columns represent all five training runs with each column label signifying the test set. Rows display the model performance for all five data sets with the training cases on the off-diagonal and the test cases on the diagonal.

Figures 9 and 10 suggest that the performance of both constitutive recurrent neural network models on the training set lies, as in the train on one task, between the performance of the neo Hookean standard linear solid model and that of the vanilla recurrent neural network. Similar to these two, the training error for the principal-stretch and invariant-based networks increases for the train on four task compared to the train on one task (increasing average NRMSE: from 0.048 to 0.208 and from 0.022 to 0.108). Looking at the test cases, the constitutive network models exhibit less overfitting compared to the vanilla network (decreasing average NRMSE: from 0.289 to 0.196 and from 0.243 to 0.159). Comparison of the train and test errors for the constitutive network models reveals that the errors are similar between train and test sets. This provides evidence that the constitutive network models experience less overfitting compared to the vanilla network for which the train and test errors show a larger difference.

**Figure 9:**
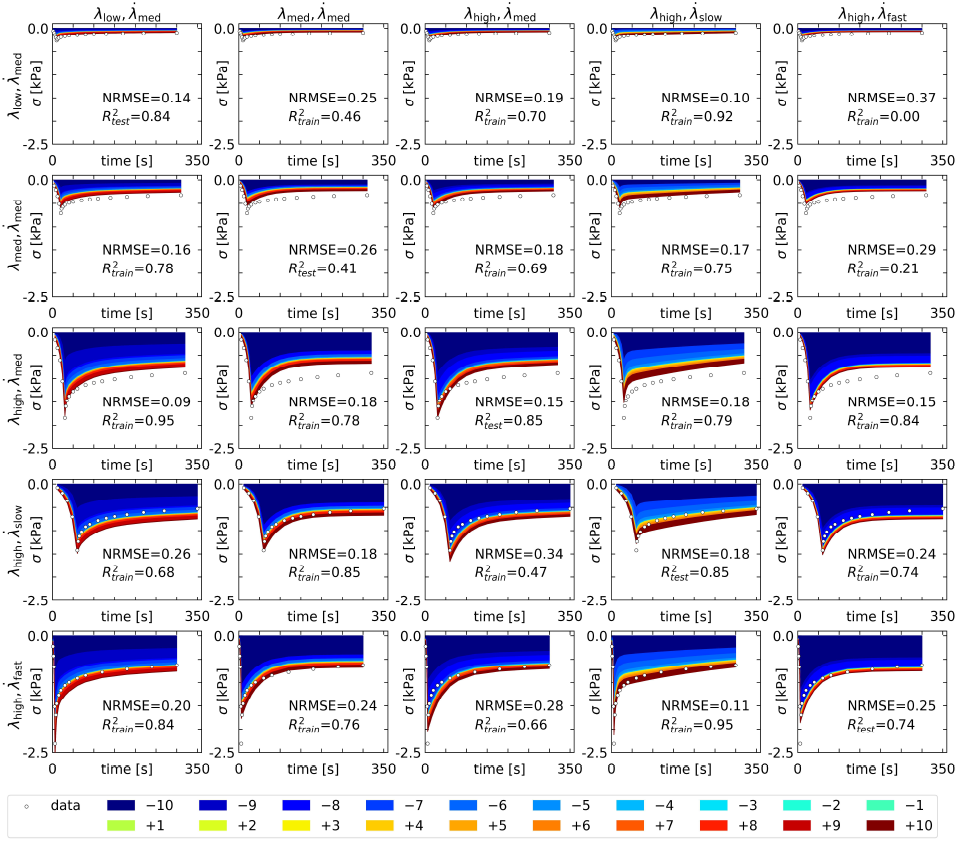
Performance of principal-stretch-based recurrent neural network trained on four curves. Experimental data and discovered model for the constitutive behavior of muscle tissue after discovering the model parameters for four of the data sets combined. Columns represent all five training runs with each column label signifying the test set. Rows display the model performance for all five data sets with the training cases on the off-diagonal and the test cases on the diagonal. The contributions of the principal stretch terms in the initial stored energy function are illustrated in colors.

**Figure 10:**
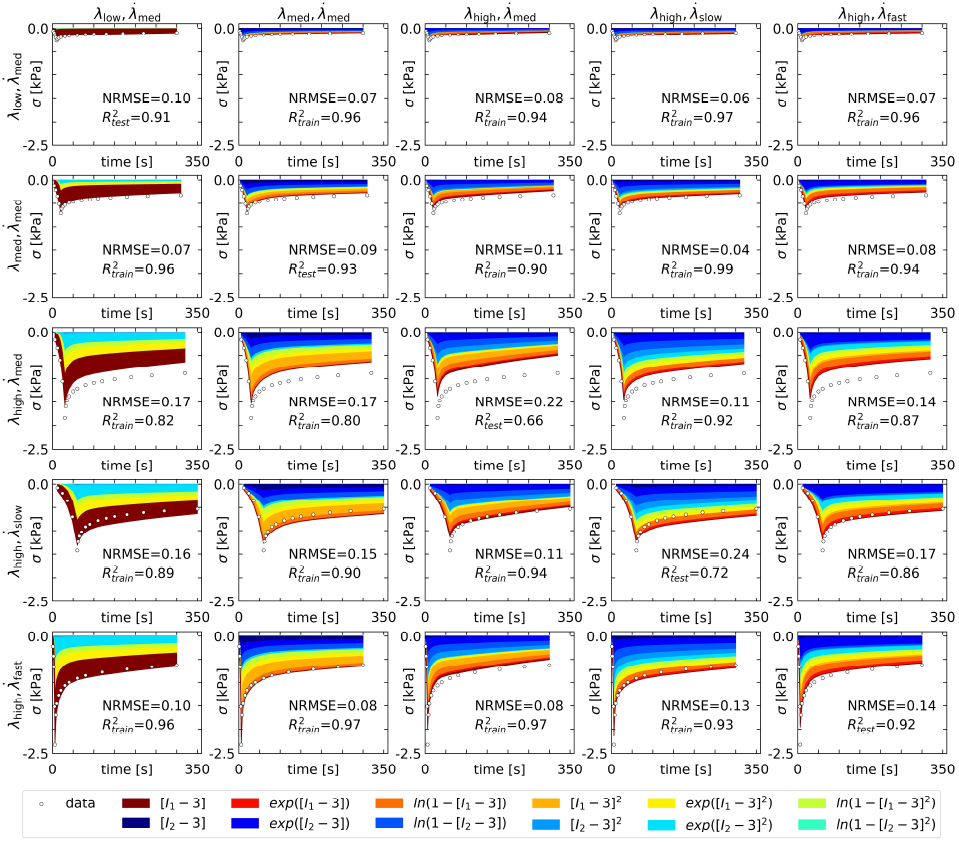
Performance of invariant-based recurrent neural network trained on four curves. Experimental data and discovered model for the constitutive behavior of muscle tissue after discovering the model parameters for four of the data sets combined. Columns represent all five training runs with each column label signifying the test set. Rows display the model performance for all five data sets with the training cases on the off-diagonal and the test cases on the diagonal. The contributions of the principal stretch terms in the initial stored energy function are illustrated in colors.

### 4.3 Model discovery

Figure 11 summarizes the performance of all four models on both evaluation tasks. The left column corresponds to the first evaluation task with training on just one curve, and the right column corresponds to the second task with training on four of the five curves. The first row displays the training errors and the second row the test errors. All displayed NRMSE values are the averages per curve in the training or test set.

**Figure 11:**
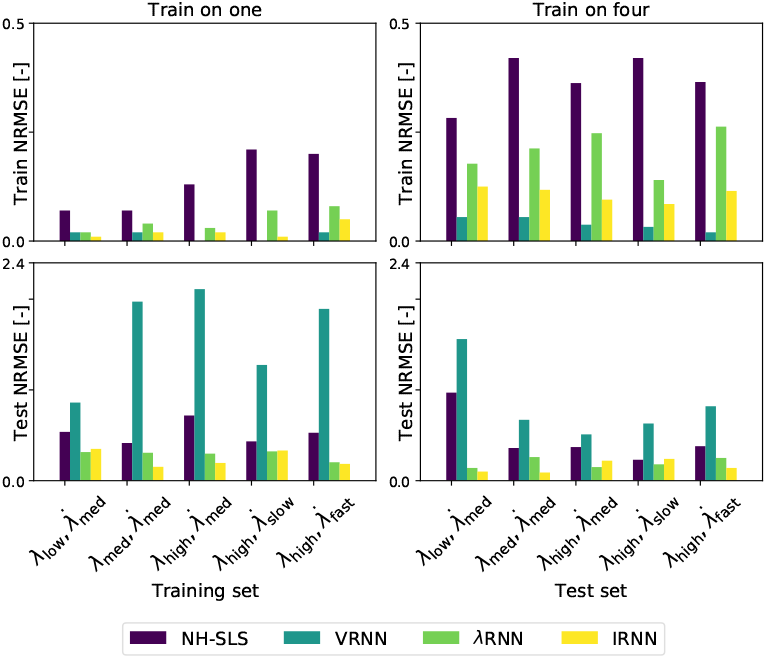
Comparison of all four models for training and testing on one and four curves. Normalized root mean squared error (NRMSE) for the train on one and train on four tasks for all four models, neo Hookean standard linear solid (NH-SLS), vanilla recurrent neural network (VRNN), principal-stretch-based (*λ*RNN) and invariant-based (IRNN) recurrent neural networks. The plotted values are the average NRMSE per curve in a given test or training set. Some bars are invisible in the top row because the errors are equal or very close to zero.

The same general trend appears in all four models where the training errors are lower than the test errors. This is expected as the models are fit to the training data and have no prior knowledge of the test data. Moving from the train on one task to the train on four task, the training errors increase as the models face the more challenging task of representing four curves using just one set of parameters. With the additional training data, however, the test errors are generally lower for the train on four case compared to the train on one case.

Focusing in on the performance of individual models, the vanilla recurrent neural network clearly fits the training data the best with errors equal to or close to zero. However, it also exhibits the largest test errors, highlighting an issue with overfitting. Both the neo Hookean standard linear solid and the constitutive recurrent neural network models exhibit less overfitting, as their train and test errors are closer in magnitude. Both constitutive recurrent neural networks outperform the neo Hookean standard linear solid overall with the invariant-based version seemingly fitting the data slightly better than the principal-stretch-based model.

### 4.4 Parameter discovery

The constitutive recurrent neural network models discover two sets of weights: one corresponding to the parameters of the initial stored energy function and one corresponding to the parameters of the Prony series relaxation function. The distribution of the discovered parameters for the initial stored energy functions is displayed using color coding in Figures 5 and 9 for the principal-stretch-based model and Figures 6 and 10 for the invariant-based model. In these figures, we assign each term in the initial stored energy function a color. The thickness of each color band indicates the relative contribution of that term to the final initial stored energy function.

Looking at the principal-stretch-based model in Figures 5 and 9, in each training scenario, the model learns different combinations of terms. However, the parameter distributions all share some general trends. For both the train on one and train on four tasks, the discovered models show a preference for negative exponent terms in the initial stored energy function. Similarly, all of the results show a preference for terms at either extreme, large negative and large positive exponents. In a study fitting a single-term Ogden model to muscle data, the resulting exponent was 14.00 for bovine and porcine tissue and 8.97 for human tissue [40]. Similarly, a study fitting a two-term compressible Ogden model found exponents of 11.77 and 14.34 for tensile data measured on guinea pig ventricular papillary muscle [18]. While our discovered models do not exactly match these fitted functional forms, our discovered exponents agree well with these large exponents reported in the literature. This suggests that the trend of our network to discover terms at either extreme agrees well with reported observations for muscle tissue.

The invariant-based model in Figures 6 and 10 behaves similarly, learning different combinations of terms for different training scenarios. However, the invariant-based model does not show any strong preference towards terms involving the first or second invariant. The model similarly does not show any strong preference towards the linear, quadratic, exponential, or logarithmic terms. Section 4.5 discusses the terms learned by the invariant-based model in more detail.

Figure 12 illustrates the distributions of the Prony parameters discovered by both constitutive recurrent neural networks. The constitutive recurrent neural network with *n*_prn_=10 in equation (32) discovers ten Prony terms, each with a corresponding time constant, *τ*_*i*_, and weight, *γ*_*i*_. Each bar in Figure 12 represents one of these terms with its horizontal location corresponding to *τ*_*i*_ and its height corresponding to *γ*_*i*_. The presence of multiple bars in the same location is indicated by a darker color since the bars are not fully opaque. The long-term modulus, *γ*_∞_, appears as a single green bar to the very left of each graph.

**Figure 12:**
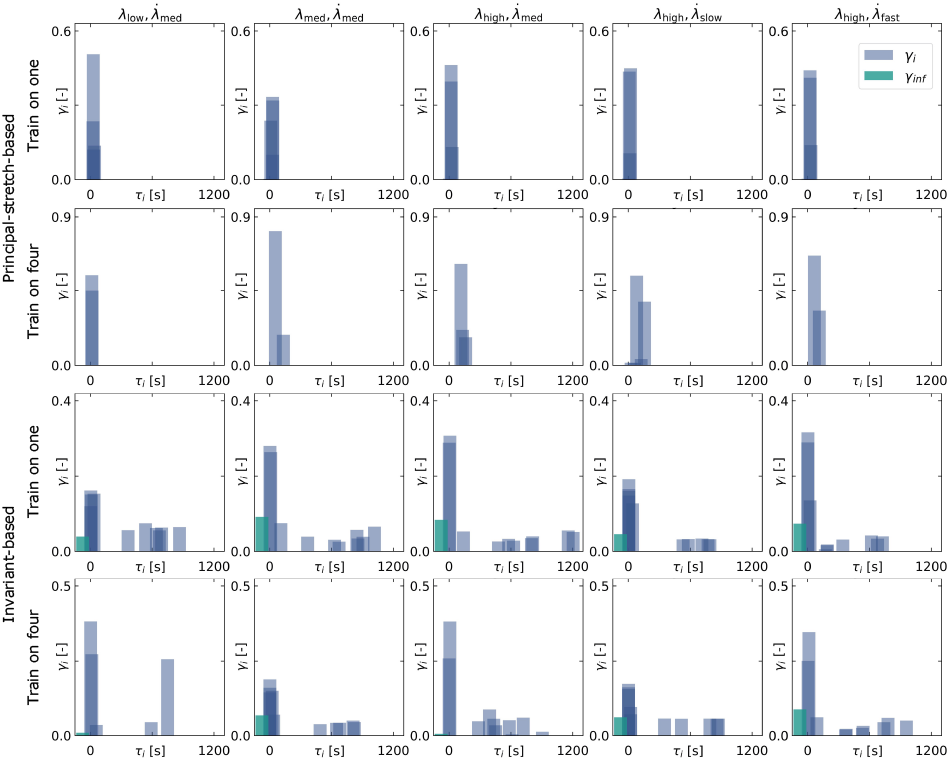
Comparison of Prony parameters for principal-stretch and invariant-based recurrent neural networks. Each bar corresponds to a time constant, *τ*_*i*_, and its height represents the magnitude of the corresponding weight, *γ*_*i*_. The bars are not fully opaque to aid the visualization of multiple terms located at similar horizontal axis locations. The long-term modulus, *γ*_∞_, is represented by a single green bar to the far left; however, it is invisible in some plots where its discovered value is close to zero.

For the principal-stretch-based model, the discovered time constants, *τ*_*i*_, in the train on one task range from 4.3s to 41s with the majority of terms concentrated around 21s to 30s. In the train on four task, the learned Prony parameters show a slightly wider spread of *τ*_*i*_ values ranging from 13s to 161s. In both the train on one and train on four cases, the principal-stretch-based model shows little contribution from the long term modulus, *γ*_∞_.

For the invariant-based model, the network discovered a greater range of *τ*_*i*_ values compared to the principal-stretch-based model. The results for both the train on one and train on four tasks look similar with *τ*_*i*_ values ranging from 1.16s to 1205s and 1.77s to 959s, respectfully. The discovered parameters for both tasks display large contributions from time constants clustered around 1s to 4s, 8s to 13s, and 22s to 40s. These main clusters are accompanied by scattered contributions from larger time constants in the range of 100s to 1200s. In contrast to the principal-stretch-based model, the invariant-based model also predicts a more significant contribution of the long term modulus, *γ*_∞_.

In a study fitting a five-term Prony series to the same muscle data [59], the resulting *τ*_*i*_ values were 0.6s, 6s, 30s, 60s, and 300s with the two smaller time constants weighted more heavily than the larger three, with corresponding weights of 0.465, 0.200, 0.057, 0.066, and 0.089. Another study fitting a two-term Prony series to guinea pig ventricular muscle data found time constants of 1.74s and 52.16s [18]. The general trends of these findings match the distributions discovered in this study with heavily-weighted contributions from time constants on the order of 1s to 10s accompanied by lesser-weighted contributions from time constants on the order of 100s.

### 4.5 Regularization

Looking at the color distributions in Figures 5, 6, 9, and 10, we recognize that both constitutive recurrent neural networks discover a broad spectrum of terms for the initial stored energy function, rather than focusing on a select few. Towards the goal of preventing potential overfitting, we investigated the ability of L2 regularization to reduce the number of discovered terms for the invariant-based model, both for the elastic and viscous parts.

First, we varied the regularization parameter *α* in equation (38) from 10^−7^ to 10^−1^. For each *α* value, we simultaneously trained the invariant-based model on all five data sets. Figure 13 displays the results for the case with no regularization in the leftmost column and cases with a gradually increasing regularization parameter *α* from left to right. From the color spectrum in Figure 13, we conclude that, as the regularization parameter *α* increases from zero to 10^−1^, the number of terms in the initial stored energy function decreases from six to two. This controlled reduction of terms agrees with the results of previous studies on the effects of regularization in constitutive neural networks for time-independent hyperelasticity [31, 51]. With no regularization, our invariant-based network discovers a six-term initial stored energy function of the following form,

**Figure 13:**
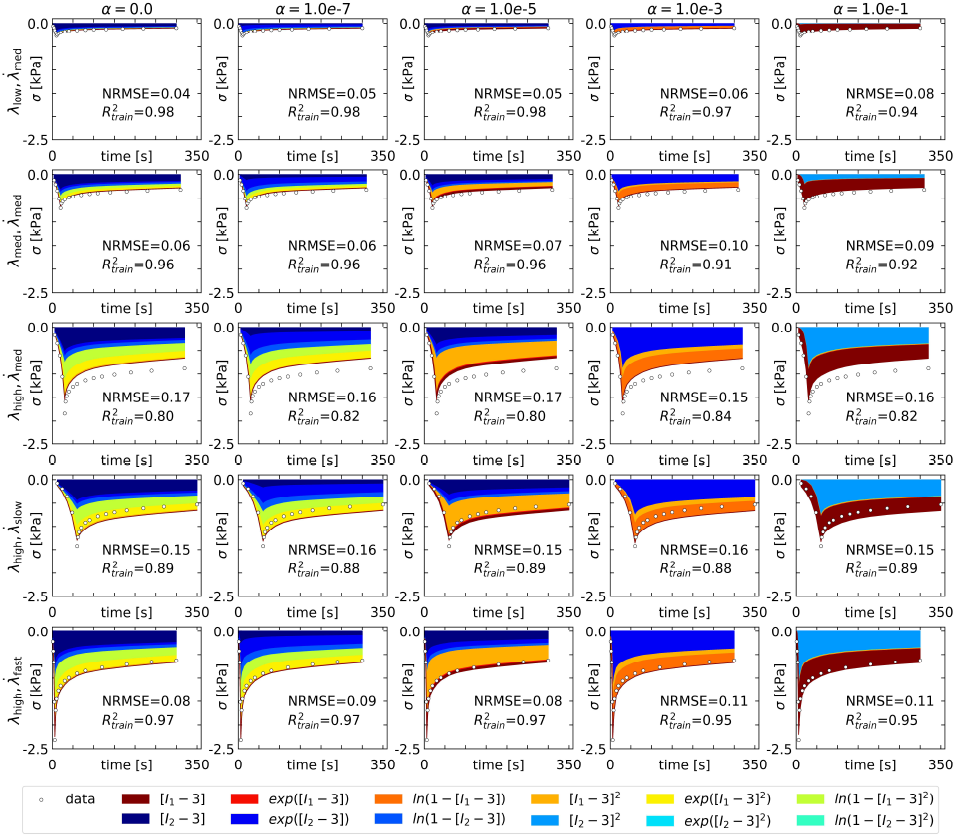
Effect of L2 regularization on initial stored energy of invariant-based neural network. Displayed results from simultaneously fitting the invariant-based recurrent neural network to all five data sets for varying regularization parameters *α*. Increasing the regularization parameter from zero via 10^−7^ to 10^−1^ decreases the number of discovered terms of the initial stored energy function from six to two. At *α* = 10^−1^, the network discovers a two-term model that is linear in the first invariant *I*_1_ and quadratic in the second invariant *I*_2_.

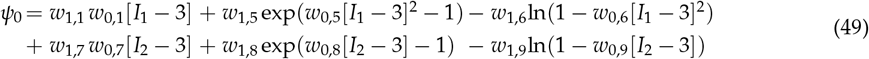

with the following weights, *w*_0,4_ = 0.19, *w*_1,4_ = 0.30kPa, *w*_0,5_ = 0.12, *w*_1,5_ = 3.69kPa, *w*_0,6_ = 0.13, *w*_1,6_ = 4.41kPa, *w*_0,7_ = 1.71, *w*_1,7_ = 0.23kPa, *w*_0,8_ = 0.11, *w*_1,8_ = 0.68kPa, *w*_0,9_ = 0.10, and *w*_1,9_ = 1.07kPa. Notably, all other weights in equation (27) train to zero. With the maximal regularization parameter of *α* = 10^−1^, our invariant-based network discovers a two-term initial stored energy function of the following form,

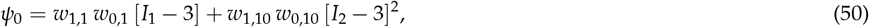

where *w*_0,1_ = 19.97kPa, *w*_1,1_ = 0.03, *w*_0,10_ = 18.37kPa, and *w*_1,10_ = 0.03, while all remaining terms train to zero. Strikingly, this final reduced version of the discovered initial stored energy function in equation (50), with a linear term in the first invariant *I*_1_ and a quadratic term in the second invariant *I*_2_, belongs to the family of generalized Mooney-Rivlin models [41, 48], 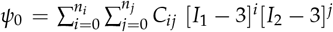, and its discovered weights translate into the Mooney-Rivlin coefficients *C*_10_ = *w*_1,1_ *w*_0,1_ = 0.60kPa and *C*_02_ = *w*_1,10_ *w*_0,10_ = 0.55kPa, with all other coefficients, *C*_*ij*_, equal to zero. In a study using the Mooney-Rivlin model with considerations for intramuscular pressure [64], the resulting coefficients were *C*_10_ = 0.05kPa and *C*_01_ = 0.50kPa for the rabbit tibialis anterior muscle. These values were obtained for modeling hyperelasticity rather than for use in modeling the inital stress response of a viscoelastic model, so direct comparison to our values is not possible. However, these values from the literature suggest that our results are of a reasonable order of magnitude.

Second, we investigated the ability of L2 regularization to reduce the number of discovered terms in the Prony series relaxation function of the invariant-based recurrent neural network. We varied the regularization parameter *β* in equation (39) from 10^−4^ to 10^−1^. For each *β* value, we simultaneously trained the invariant-based model on all five data curves. Figure 14 displays the results for the case with no regularization in the leftmost column and cases with a gradually increasing regularization parameter *β* from left to right. The top row displays the distribution of learned Prony parameters in the same format as Figure 12, and the remaining rows show the model predictions on the five curves. Looking at the top row of Figure 14, as we increase the regularization parameter *β* from zero to 10^−1^, the number of learned Prony terms decreases from seven to two. However, for the largest value of *β* = 10^−1^, the errors between the model predictions and the experimental data increase significantly. This suggests that the optimal regularization parameter lies below this value, *β <* 10^−1^. With no regularization, the invariant-based network discovers a seven-term Prony series for the relaxation function:

**Figure 14:**
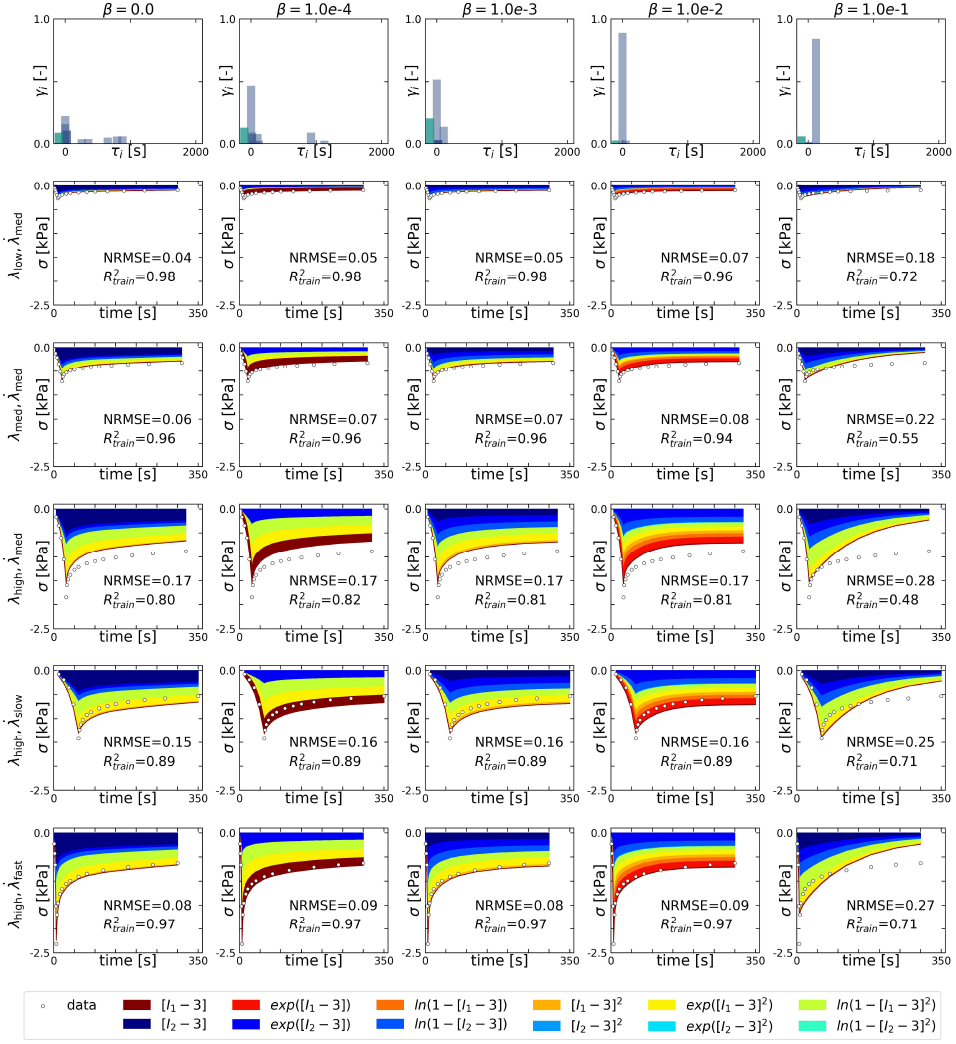
Effect of L2 regularization on Prony parameters of recurrent neural network. Displayed results from simultaneously fitting the invariant-based recurrent neural network to all five data sets for varying regularization parameters *β*. Increasing the regularization parameter from zero via 10^−4^ to 10^−1^ decreases the number of discovered terms of the Prony series from six to two. At *β* = 10^−2^, the network discovers a three-term Prony series with time constants of 0.362s, 2.54s, and 52.0s.

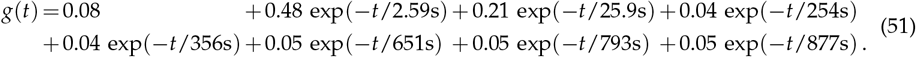

As we increase the regularization parameter *β*, the model discovers fewer terms; yet, at the same time, the prediction error increases significantly. A value of *β* = 10^−2^ for which the network discovers a three-term relaxation function,

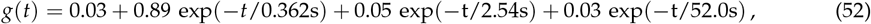

seems to be a reasonable trade off between model terms and model error. The magnitudes of the three time constants discovered by the regularized model closely match the time constants of 1.74s and 52.16s reported for a two-term Prony series fitted to guinea pig ventricular muscle data [18]. Our three discovered time constants are also comparable to the range of time constants, from 0.6s-300s, used to fit the same muscle data to a five-term Prony series [59]. Our discovered time constants lie in the lower end of the 0.6-300s range, and we note that the L2 regularization trends towards smaller time constants for increasing *β* as seen in Figure 14.

## 5 Limitations and future outlook

While our two constitutive recurrent neural networks in their principal-stretch-based and invariant-based versions show promise in the discovery of constitutive models for muscle tissue, our current model also has some limitations to address: First, as a proof of concept, we have focused on a very specific case of uniaxial unconfined compression and have developed a one-dimensional model in accordance with this loading configuration. Second, for simplicity, we only consider incompressibility and isotropy. Moving forward, we will expand our general framework to accommodate other loading scenarios by extending it to three dimensions [9, 31] and incorporating considerations for compressibility [18, 64] and anisotropy [4, 25, 26, 29, 52, 53, 58]. A three-dimensional formulation would allow the investigation of different loading modes such tension, compression, and shear [5] or biaxial stretch [62] and the investigation of tension-compression asymmetry [27, 51]. For hyperelastic materials, we have shown that including different loading modes in the training process is critical to reproducibly discover a single unique model [31], and we are confident that this stabilizing effect will translate to viscoelastic model discovery. Third, our model is limited by the assumptions of the theory of quasi-linear viscoelasticity [13]. Following this theory, our model assumes that the material response can be decomposed into a time-independent initial stress function that depends only on the current stretch and a time-dependent relaxation function that depends only on time. The theory of quasi-linear viscoelasticity is widely used to model single cells [61] and tissues, such as tendons [45, 66], skeletal muscle [59], or the heart [57]. Materials for which this assumption does not hold will require a different, fully-nonlinear modeling approach [Tac et al., 2023b, 1, 20, 28, 63, 65]. However, as a reasonable first approximation, the theory of quasi-linear viscoelasticity characterizes the behavior of muscle tissue well [18, 40, 47, 56, 59, 64] and produces comparable results in this study.

## 6 Conclusions

Constitutive artificial neural networks are pioneering a new approach to constitutive modeling where both model and parameters are fit to the data themselves. Our results show that a recurrent neural network architecture extends the capabilities of past feed-forward networks to model the history-dependent behavior of viscoelastic materials. Inspired by the theory of quasi-linear viscoelasticity, we translate our domain knowledge into a new class of recurrent neural networks that requires less training data than classical vanilla recurrent neural networks. Our results show that given limited training data from mechanical testing of muscle tissue, a vanilla recurrent neural network struggles to predict stress responses for unseen data, often generating predictions that violate basic physical principles. In contrast, our constitutive recurrent neural network combines the advantages of an overly-constrained neo Hookean standard linear solid and an overly-flexible vanilla recurrent neural network. It robustly discovers constitutive models that obey physical principles and generalize well to unseen data. Furthermore, the modular structure of our network architecture takes advantage of past and ongoing research on hyperelastic feed-forward networks by extending these architectures with an additional recurrent neural network layer. This modular structure naturally encourages the plug-and-play of different families of hyperelastic models– principal-stretch or invariant-based, isotropic or anisotropic, incompressible or compressible–and shapes the path for future studies to discover relaxation functions by providing a clear map for model implementation.

## Credit authorship contribution statement

Lucy Wang: conceptualization, methodology, investigation, visualization, writing - original draft, writing - review & editing; Kevin Linka: methodology, writing - review & editing; Ellen Kuhl: conceptualization, supervision, writing - original draft, writing - review & editing

## Data availability

Our code, data, and examples are available at https://github.com/LivingMatterLab.

